# Clonal hematopoiesis in individuals with *ANKRD26* or *ETV6* germline mutations

**DOI:** 10.1101/2021.11.17.468983

**Authors:** Michael W. Drazer, Claire C. Homan, Kai Yu, Marcela Cavalcante de Andrade Silva, Kelsey E. McNeely, Matthew J. Pozsgai, Maria G. Acevedo, Jeremy P. Segal, Peng Wang, Jinghua Feng, Sarah L. King-Smith, Erika Kim, Sophia C. Korotev, David M. Lawrence, Andreas W. Schreiber, Christopher N. Hahn, Hamish S. Scott, Raman Sood, NISC Comparative Sequencing Program, Elvira D R P Velloso, Anna L. Brown, Paul P. Liu, Lucy A. Godley

## Abstract

Currently, there are at least a dozen recognized hereditary hematopoietic malignancies (HHMs), some of which phenocopy others. Among these, three HHMs driven by germline mutations in *ANKRD26, ETV6*, or *RUNX1* share a phenotype of thrombocytopenia, qualitative platelet defects, and an increased lifetime risk of hematopoietic malignancies (HMs). Prior work has demonstrated that *RUNX1* germline mutation carriers experience an elevated lifetime risk (66%) for developing clonal hematopoiesis (CH) prior to age 50. Germline mutations in *ANKRD26* or *ETV6* phenocopy *RUNX1* germline mutations, but no studies have focused on the risk of CH in individuals with germline mutations in *ANKRD26* or *ETV6*.

To determine the prevalence of CH in individuals with germline mutations in *ANKRD26* or *ETV6*, we performed next generation sequencing on hematopoietic tissue from twelve individuals with either germline *ANKRD26* or germline *ETV6* mutations. Each patient had thrombocytopenia but had not developed HMs. Among the seven individuals with germline *ANKRD26* mutations, one patient had a CH clone driven by a somatic *SF3B1* mutation (p.Lys700Glu). This mutation increased from a variant allele frequency (VAF) of 9.4% at age 56 to 17.4% at age 60. None of the germline *ETV6* mutation carriers had evidence of CH at the limits of detection of the NGS assay (5% VAF). Unlike individuals with germline mutations in *RUNX1*, no individuals under the age of 50 with germline mutations in *ANKRD26* or *ETV6* had detectable CH. This work demonstrates that *ANKRD26* germline mutation carriers, but not *ETV6* mutation carriers, experience elevated risk for CH.

## To the Editor

Currently, there are at least a dozen recognized hereditary hematopoietic malignancies (HHMs), some of which phenocopy others. Among these, three HHMs driven by germline mutations in *ANKRD26, ETV6*, or *RUNX1* share a phenotype of thrombocytopenia, qualitative platelet defects, and an increased lifetime risk of hematopoietic malignancies (HMs).^1^ Individuals with germline mutations in these hereditary thrombocytopenia/hereditary hematopoietic malignancy (HT/HHM) associated genes experience a lifetime risk for HMs of approximately 8% (*ANKRD26*), 33% (*ETV6*), or 44% (*RUNX1*).^1^

Nine unaffected *RUNX1* germline mutation carriers with thrombocytopenia, but no HMs, were sequenced in a previous study. This demonstrated that 66% of these individuals had clonal hematopoiesis (CH) prior to age 50, an elevated CH risk as compared to population controls.^2, 3^ A subsequent study of four *RUNX1* germline mutation carriers (age 49, 53, 56, and 71 years) with thrombocytopenia, but no HMs, demonstrated CH in three of these individuals.^4^ Germline mutations in *ANKRD26* or *ETV6* phenocopy *RUNX1* germline mutations, but no studies have focused on the risk of CH in individuals with germline mutations in *ANKRD26* or *ETV6*.

To address this knowledge gap, we performed a cross sectional study of twelve individuals with either germline *ANKRD26* or germline *ETV6* mutations who had thrombocytopenia but who had not developed HMs. We determined if *ANKRD26* or *ETV6* germline mutations lead to increased rates of CH, as is observed in *RUNX1* mutation carriers.^2, 4^ Given that the penetrance of HMs is lower in *ANKRD26* and *ETV6* germline mutation carriers than in *RUNX1* mutation carriers, we hypothesized that germline *ANKRD26* or *ETV6* mutation carriers would experience lower rates of CH relative to germline *RUNX1* mutation carriers of similar ages.^1^ Additionally, all pathogenic/likely pathogenic *ANKRD26* variants with supporting evidence in ClinVar are located in a regulatory domain, the 5’ untranslated region (UTR).^5^ *RUNX1* encodes for a transcription factor (RUNX1) that binds to the *ANKRD26* 5’ UTR and suppresses *ANKRD26* expression **(Supplementary Figure 1**).^5^ Therefore, we hypothesized that *ANKRD26* and *RUNX1* germline mutation carriers would experience somatic mutations in a similar set of genes as compared to *ETV6* germline mutation carriers.

We enrolled twelve patients from unrelated families on Institutional Review Board-approved protocols at the University of Chicago (UChicago) or the Hospital das Clínicas da Faculdade de Medicina da Universidade de São Paulo, Brazil (**Supplementary Table 1**). Seven unaffected *ANKRD26* mutation carriers and four unaffected *ETV6* mutation carriers were enrolled. Patient ages ranged from 8 to 63 years. Additionally, we included one affected *ANKRD26* mutation carrier with acute myeloid leukemia (AML) with 23% blasts in order to compare CH-related mutations to an HM in this syndrome (**Table 1, Supplementary Table 1**). **Supplementary Figure 2** shows a pedigree for each family. Each HT/HHM-related germline variant was classified using Association for Molecular Pathology and American College of Human Genetics and Genomics criteria.^6^ Patient samples included germline tissue (cultured skin fibroblasts) or hematopoietic tissue equivalents (peripheral blood, bone marrow, or saliva). Panel-based sequencing was performed at UChicago as described previously.^7^ Details regarding sample processing and sequencing are in the **Supplementary Methods** section.

**Table 1.** Germline mutations and somatic driver mutations identified in each individual in *ANKRD26* and *ETV6*-mutated HT/HHM phenocopy cohort. Family numbers and Individual IDs reference pedigrees shown in Supplementary Figure 2.

| **Unaffected Individuals** |  |  |  |  |  |  |  |
| --- | --- | --- | --- | --- | --- | --- | --- |
| **Germline Gene** | **Variant** | **Family** | **Individual ID** | **Age** | **CH present?** | **CH mutation** | **VAF** |
| *ANKRD26* | 5' UTR c.-128G>A NM_014915.2 | 4 | III-16 | 8 | N | NA | NA |
| *ANKRD26* | 5' UTR c.-128G>A NM_014915.2 | 4 | III-14 | 22 | N | NA | NA |
| *ANKRD26* | 5' UTR c.-118C>T NM_014915.2 | 2 | IV-1 | 25 | N | NA | NA |
| *ANKRD26* | 5' UTR c.-118C>T NM_014915.2 | 2 | III-4 | 55 | N | NA | NA |
| *ANKRD26* | 5' UTR c.-118C>T NM_014915.2 | 2 | III-2 | 56 | Y | *SF3B1* p.Lys700Glu | 9.4% |
| *ANKRD26* | 5' UTR c.-118C>T NM_014915.2 | 2 | III-2 | 60 | Y | *SF3B1* p.Lys700Glu | 17.4% |
| *ANKRD26* | 5' UTR c.-119C>G NM_014915.2 | 3 | IV-3 | 43 | N | NA | NA |
| *ANKRD26* | 5' UTR c.-119C>G NM_014915.2 | 3 | IV-3 | 44 | N | NA | NA |
| *ANKRD26* | 5' UTR c.-119C>G NM_014915.2 | 3 | V-1 | 13 | N | NA | NA |
| *ETV6* | p.R369Q NM_001987.4 | 1 | IV-5 | 33 | N | NA | NA |
| *ETV6* | p.R369Q NM_001987.4 | 1 | IV-4 | 36 | N | NA | NA |
| *ETV6* | p.R369Q NM_001987.4 | 1 | IV-4 | 38 | N | NA | NA |
| *ETV6* | p.R369Q NM_001987.4 | 1 | III-3 | 59 | N | NA | NA |
| *ETV6* | p.R369Q NM_001987.4 | 1 | III-3 | 62 | N | NA | NA |
| *ETV6* | p.R369Q NM_001987.4 | 1 | III-5 | 62 | N | NA | NA |
| *ETV6* | p.R369Q NM_001987.4 | 1 | III-3 | 63 | N | NA | NA |
| **Affected Individuals** |  |  |  |  |  |  |  |
| **Germline Gene** | **Variant** |  | **Individual ID** | **Age** | **Malignancy** | **Somatic Driver Mutations** | **VAF** |
| *ANKRD26* | 5' UTR c.-128G>A NM_014915.2 | 4 | II-12 | 48 | AML (23% blasts) | *CUX1* p. Phe472GlnfsX105 | 92.4% |
|  |  |  |  |  |  | *RUNX1* p.Arg320X | 55.8% |
|  |  |  |  |  |  | *TET2* p.Phe1309LeufsX54 | 40.0% |
|  |  |  |  |  |  | *FLT3* p.Asp835His | 18.3% |
|  |  |  |  |  |  | *SAMD9* p.Val798GlyfsX7 | 13.3% |

The ages of individuals with germline *ANKRD26* or *ETV6* mutations at the time of sample collection are shown in **Table 1**. The median age at sample collection was 43 years for unaffected *ANKRD26* germline mutation carriers, with serial samples from one individual. The median age for unaffected *ETV6* germline mutation carriers was 59 years, with serial samples from two individuals.

Among the seven individuals with germline *ANKRD26* mutations, one patient with a germline *ANKRD26* mutation had a CH clone driven by a somatic *SF3B1* mutation (p.Lys700Glu). This mutation increased from a variant allele frequency (VAF) of 9.4% at age 56 to 17.4% at age 60 (**Table 1, Figure 1**). *SF3B1* p.Lys700Glu is a recognized somatic hotspot mutation that is observed in 2.1% of COSMIC HMs (n = 525/25028).^8^ Of note, the only patient with CH in the *ANKRD26* cohort was also the oldest individual in that cohort. None of the germline *ETV6* mutation carriers (n = 4) had evidence of CH at the limits of detection of the NGS assay (5% VAF). Unlike individuals with germline mutations in *RUNX1*, no individuals under the age of 50 with germline mutations in *ANKRD26* (n=6) or *ETV6* (n=3) had detectable CH despite nearly half of the unaffected samples being collected from individuals in this age group (**Figure 1**).^2, 4^

**Figure 1.**
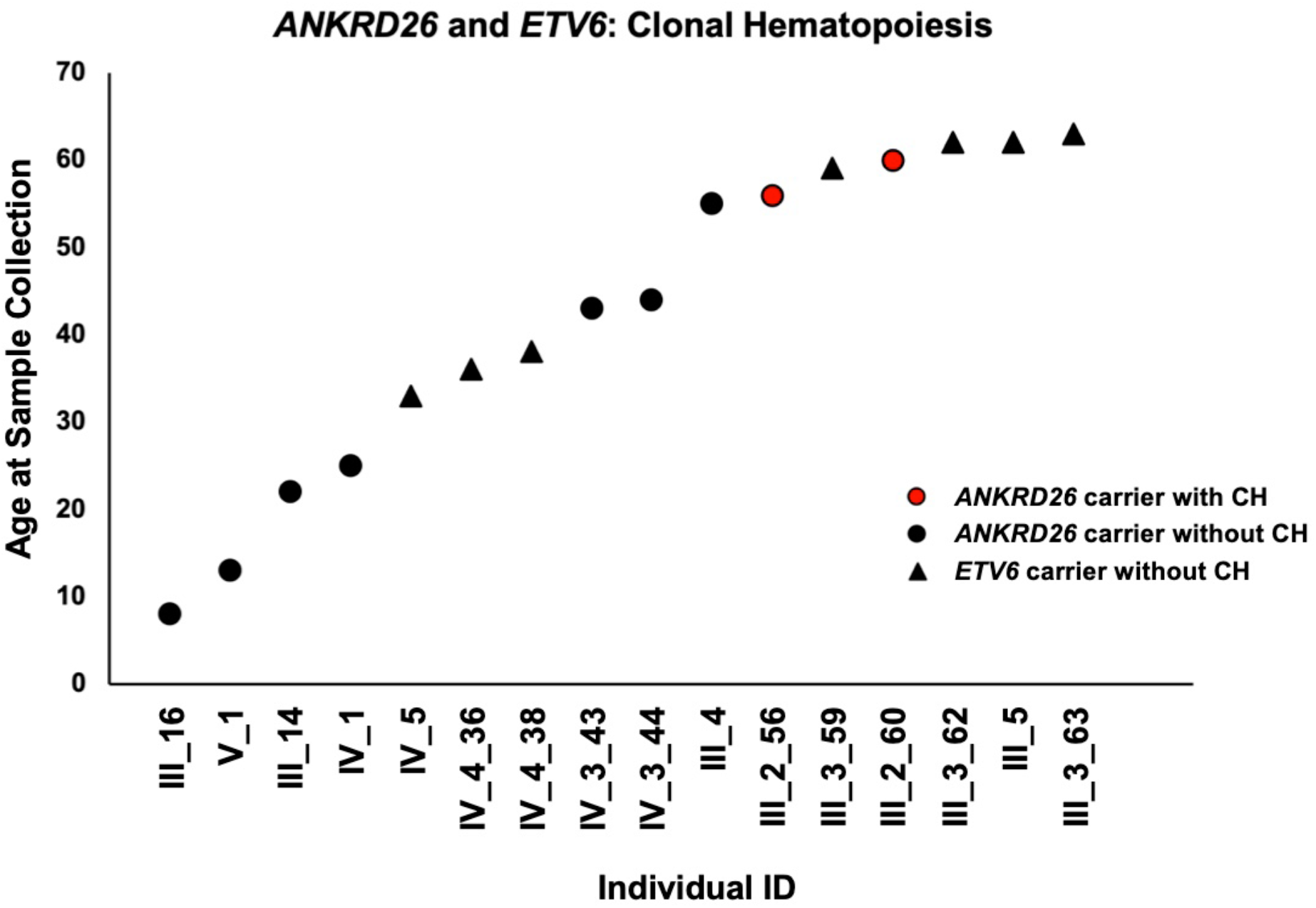
Clonal hematopoiesis in *ANKRD26* or *ETV6* germline mutation carriers. One individual with a germline *ANKRD26* mutation (5’ UTR, c.-118C>T, NM_014915.2) had a CH clone driven by *SF3B1* p.Lys700Glu in two samples collected at the ages of 56 and 60.

The only patient with a germline *ANKRD26* mutation and a malignancy (AML) had a leukemic clone with both typical and atypical driver mutations: *CUX1* (p. Phe472GlnfsX105, VAF 92.4%), *RUNX1* (p.Arg320X, VAF 55.8%), *TET2* (p.Phe1309LeufsX54, VAF 40.0%), *FLT3* (p.Asp835His, second tyrosine kinase domain (TKD), VAF 18.3%), and *SAMD9* (p.Val798GlyfsX7, VAF 13.3%) (**Table 1**). *FLT3* is mutated in 18.8% of COSMIC HMs,^8^ and *RUNX1* somatic mutations are the most common second hit in *RUNX1* germline mutation carriers who have developed HMs.^8, 9^ Given the role of RUNX1 in regulating *ANKRD26* expression, the *RUNX1* mutation in this *ANKRD26* germline mutation carrier may effectively represent a second hit that serves as a late leukemogenic event.^5^ The *CUX1* and *SAMD9* mutations were not described previously in COSMIC.^8^ The leukemic karyotype was 46, XX, -6, del(7)(q11.2),+mar[20].

The total observation time for the *ANKRD26* cohort was 275 years. The incidence of CH in the *ANKRD26* cohort was 4.5×10^−3^ CH cases/observation year (4.5 CH cases per 1000 observation years). The incidence rate of HMs in the *ANKRD26* cohort was 3.6×10^−3^ malignancies/observation year (3.6 malignancies per 1000 observation years). This HM incidence rate was similar to that seen previously in a cohort of Italian germline *ANKRD26* mutation carriers (2.13 malignancies per 1000 observation years).^10^ The observation time for the *ETV6* cohort was 258 years, with no diagnoses of CH or HMs.

Among the known HT/HHM phenocopies, only *RUNX1* has been systematically evaluated for CH risk. This bias has likely occurred for two reasons. First, *RUNX1*-driven HT/HHMs were identified 12 years before *ANKRD26*-driven HT/HHMs and 16 years before *ETV6*-driven HT/HHMs, which has provided a longer period of time for researchers to identify and study families with germline *RUNX1* mutations.^1, 11-13^ Second, *RUNX1*-driven HT/HHMs have the highest penetrance for HMs among the HT/HHM phenocopies, with approximately 44% of mutation carriers developing blood cancers. This penetrance is higher than that experienced by *ANKRD26* (8%) and *ETV6* (33%) germline mutation carriers.^1^ In our clinical experience, the most “severe” hereditary syndromes are more easily recognized than syndromes with more subtle symptoms and lower penetrance phenotypes. Therefore, it is not surprising that prior work in the HHM field has largely focused on *RUNX1*-driven HT/HHMs.

To our knowledge, this is the first study examining pre-leukemic states in the HT/HHM phenocopies driven by germline *ANKRD26* or *ETV6* mutations. In our cohort, CH was detected in 14% of *ANKRD26* germline mutation carriers, but no CH was present in *ETV6* germline mutation carriers. No *ANKRD26* or *ETV6* mutation carriers developed malignancies during 533 years of observation time. It is possible the limited number of germline variants and families in this study, with four families carrying three *ANKRD26* variants and one family with one *ETV6* variant, are not representative of the leukemogenic risk observed in the full spectrum of HT/HHM-related *ANKRD26* or *ETV6* germline variants. Ultimately, larger numbers of germline *ANKRD26* or *ETV6* mutation carriers should be studied to better determine the pre-leukemic genetic milieu that exists in these syndromes.

In conclusion, this is the first cross sectional study focused on leukemogenic mechanisms in individuals with *ANKRD26*-or *ETV6*-driven HT/HHM phenocopies. We identified CH in 14% of older germline *ANKRD26* mutation carriers but did not detect CH in *ETV6* germline mutation carriers. We did not detect early-onset CH under the age of 50 in individuals with germline *ANKRD26* or *ETV6* mutations, as has been observed in *RUNX1* mutation carriers.^2^ We also identified a rare somatic *RUNX1* mutation in a germline *ANKRD26* mutation carrier with AML, which may effectively represent a second hit event given the role of RUNX1 in regulating *ANKRD26* expression.^5^ Given the relatively small sample size of our cohort and the limited number of pedigrees with germline *ANKRD26* mutations worldwide, future studies focused on evaluating leukemogenic mutations before and after the development of HMs in *ANKRD26* or *ETV6* germline mutation carriers should be performed.

## Acknowledgements

We thank the patients and their families for their participation in this research program and for providing samples. This work was supported by the Damon Runyon Cancer Research Foundation Physician-Scientist Training Award, the Edward P. Evans Foundation Young Investigator Award, the Cancer Research Foundation Young Investigator Award, and the NIH Paul Calabresi K12 Program in Oncology (MWD). KY, EK, RS, and PPL are supported by the Intramural Research Program at the National Human Genome Research Institute, NIH.

## Supplementary Files

**Supplementary Table 1.** Cohort of patients enrolled on the *ANKRD26* and *ETV6* cross sectional study.

| Germline Gene of Interest | Variants | Individual | Family | Phenotype | Age(s) at Sample Collection (years) |
| --- | --- | --- | --- | --- | --- |
| *ETV6* | p.R369Q NM_001987.4 | III_3 | Family 1 | Thrombocytopenia | 59, 62, 63 |
| *ETV6* | p.R369Q NM_001987.4 | IV_4 | Family 1 | Thrombocytopenia | 36, 38 |
| *ETV6* | p.R369Q NM_001987.4 | IV_5 | Family 1 | Thrombocytopenia | 33 |
| *ETV6* | p.R369Q NM_001987.4 | III_5 | Family 1 | Thrombocytopenia | 62 |
| *ANKRD26* | 5' UTR c.-118C>T NM_014915.2 | III_2 | Family 2 | Thrombocytopenia | 56, 60 |
| *ANKRD26* | 5' UTR c.-118C>T NM_014915.2 | III_4 | Family 2 | Thrombocytopenia | 55 |
| *ANKRD26* | 5' UTR c.-118C>T NM_014915.2 | IV_1 | Family 2 | Thrombocytopenia | 25 |
| *ANKRD26* | 5' UTR c.-119C>G NM_014915.2 | IV_3 | Family 3 | Thrombocytopenia | 43, 44 |
| *ANKRD26* | 5' UTR c.-119C>G NM_014915.2 | V_1 | Family 3 | Thrombocytopenia | 13 |
| *ANKRD26* | 5' UTR c.-128G>A NM_014915.2 | III_14 | Family 4 | Thrombocytopenia | 22 |
| *ANKRD26* | 5' UTR c.-128G>A NM_014915.2 | II_12 | Family 4 | AML (23% blasts) | 48 |
| *ANKRD26* | 5' UTR c.-128G>A NM_014915.2 | III_16 | Family 4 | Thrombocytopenia | 8 |

**Supplementary Figure 1.**
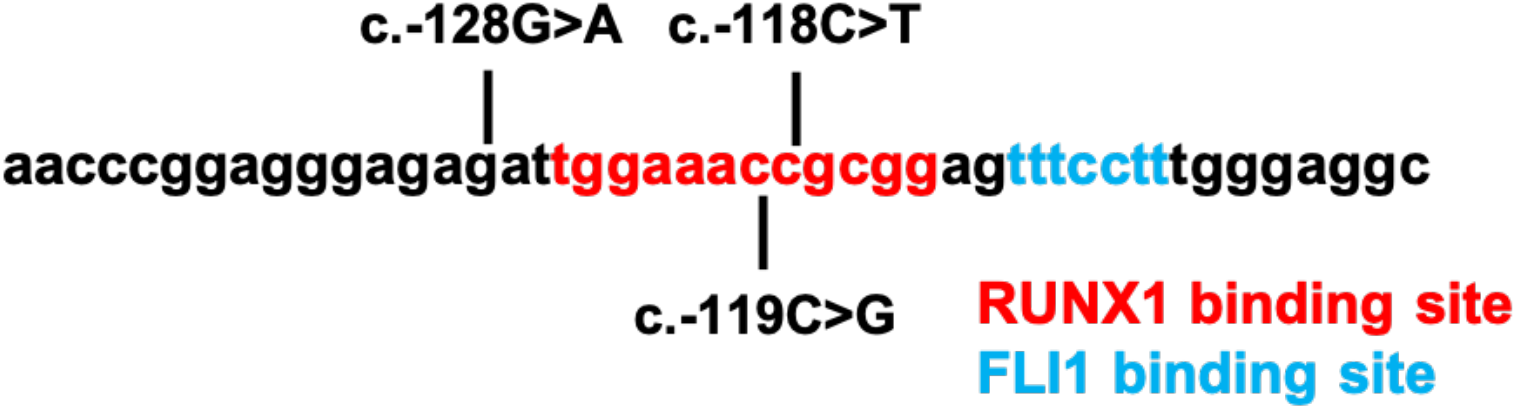
The 5’ UTR region of *ANKRD26* contains a binding site for both RUNX1 and FLI1. Germline mutations in the 5’ UTR of *ANKRD26* disrupt this interaction and lead to a hereditary thrombocytopenia/hereditary hematopoietic malignancy (HT/HHM) phenotype. This phenotype phenocopies germline *RUNX1* mutations. Germline *FLI1* mutations lead to hereditary thrombocytopenia but are not known to cause HHMs and are therefore not considered to represent an HT/HHM phenocopy. The mutations shown (c.-128G>A, c.-119C>G, and c.-118C>T) represent the mutations in the cohort of *ANKRD26* germline mutation patients in this study. ClinVar classifies the c.-118C>T and c.-119C>G variants as likely pathogenic and the c.-128G>A variant as pathogenic.

**Supplementary Figure 2.**
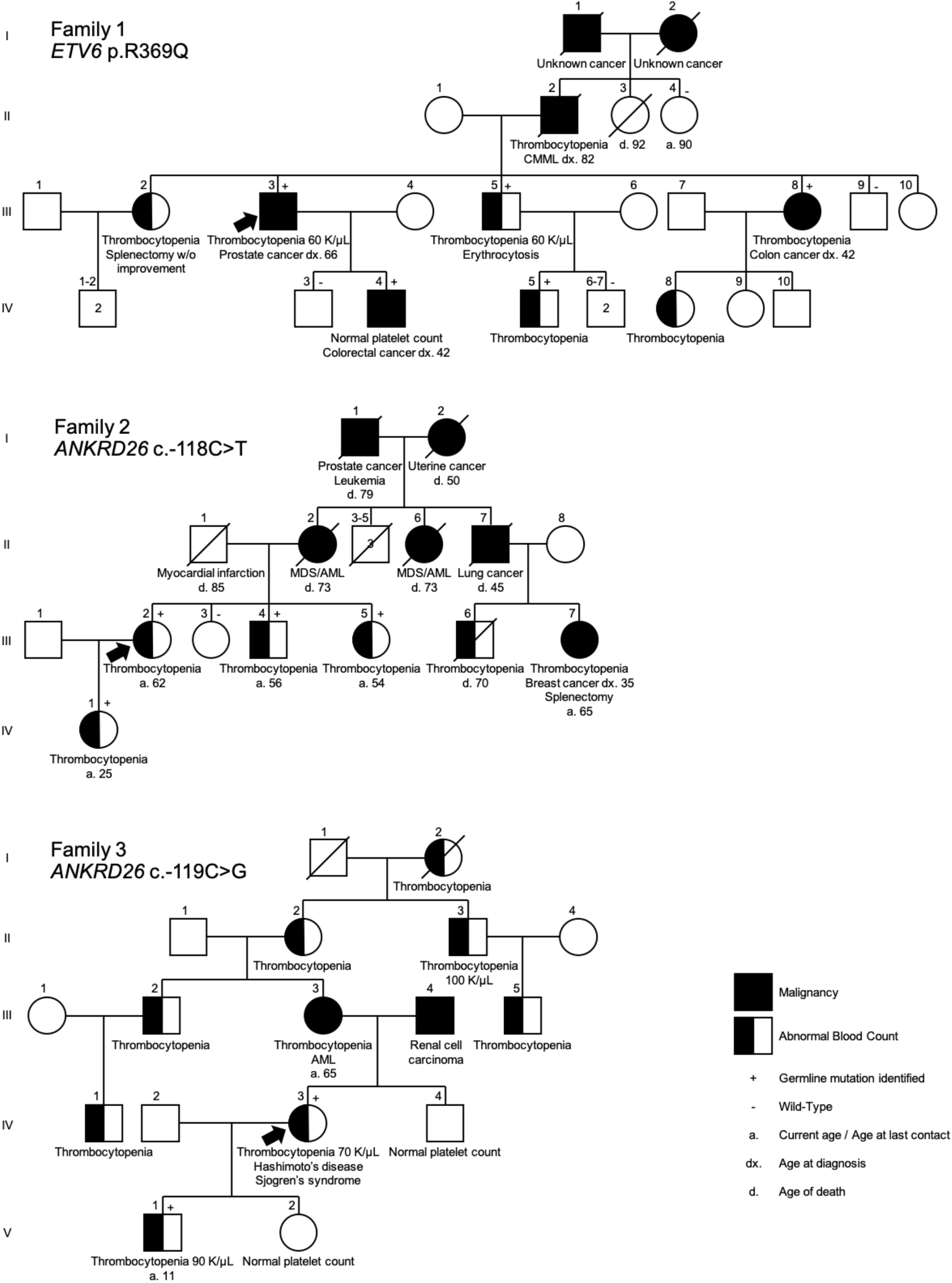
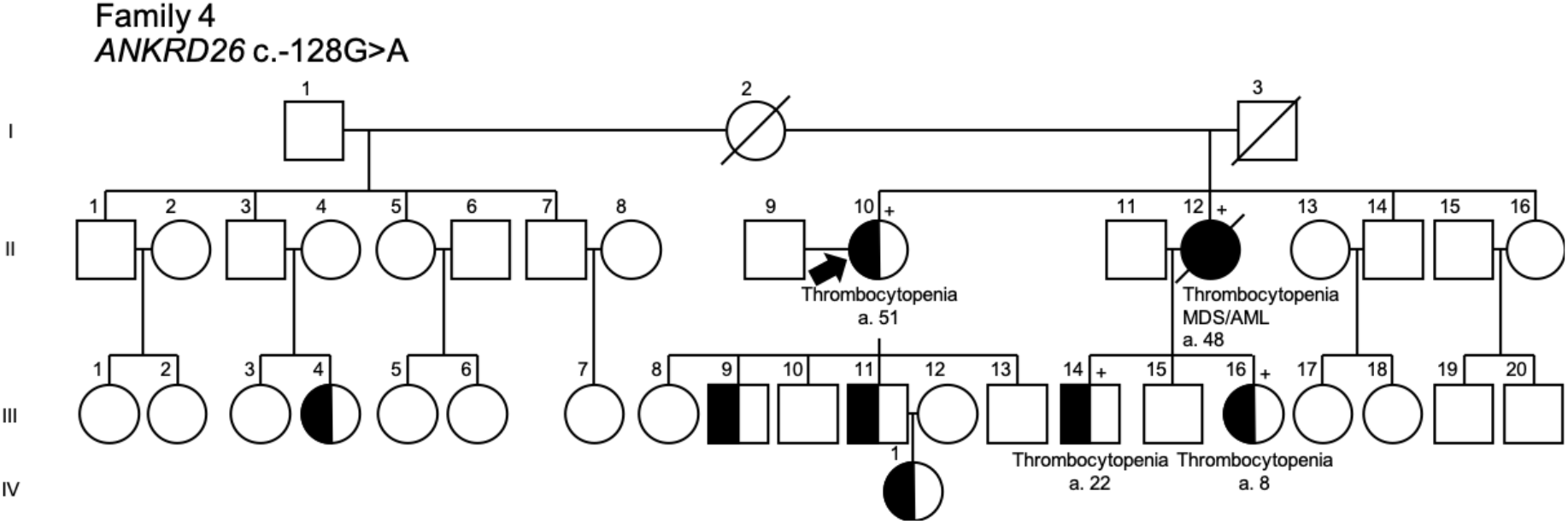
Pedigrees from families enrolled on study. *Family 1*: germline *ETV6* p.R369Q mutation family; *Family 2*: germline *ANKRD26* c.-118C>T mutation family; *Family 3*: germline *ANKRD26* c.-119C>G mutation family. Pedigree from families enrolled on study. *Family 4*: germline *ANKRD26* c.-128G>A mutation family.

## Supplementary Methods

### Sample processing and next generation sequencing methods

Genomic DNA (gDNA) was extracted with the QIAamp DNA Blood Mini Kit (Qiagen) following the manufacturer’s instructions. DNA concentrations were measured via a Nanodrop (Thermo Scientific) and/or Qubit fluorometer (Life Technologies). At least 100 ng of genomic DNA from each sample was sheared, selected by size, ligated to adapters, and standard sequencing libraries were generated via PCR amplification. Following library generation, genomic capture was performed using a custom SeqCap EZ capture panel that covered 1212 genes (Roche), and an additional PCR amplification with real-time quantitative PCR quantification was performed. An Illumina HiSeq was used to sequence the pooled capture libraries. Sequencing data were stored on a protected high-performance computing system at UChicago that exceeds requirements for the Health Insurance Portability and Accountability Act. The data were initially analyzed via a bioinformatics pipeline that melded publicly available packages built off of the GATK package and a custom bioinformatics pipeline developed at UChicago. These data were initially reviewed by MWD and KY for driver mutations.^14^ Following an initial round of review, the raw FASTQ files were then transferred to the University of South Australia in order to analyze the data using the freebayes-based RUNX1db bioinformatics pipeline.^15^ The data were filtered for read quality and depth as previously described, with thresholds as follows: variant allelic depth >= 5, read depth >= 20, population prevalence (variants at 0.1% or higher in any population database were removed), pathogenicity (missense variants that were not predicted to be damaging in 2 or more *in silico* predictors were removed; CADD scores with values less than 20 or higher were removed), and oncogenicity (variants not in genes with known roles as drivers in myeloid malignancies, not in COSMIC, or *RUNX1* variants were removed). We analyzed the subsequent list of candidate variants and used IGV to review each variant of interest manually in the individual BAM files. We removed any variants labeled as artifacts after the aforementioned steps.

We then analyzed the remaining IGV-confirmed variants to label each variant as germline or somatic in origin. For individuals with sequencing data from cultured skin fibroblasts, we compared variants identified in hematopoietic tissue equivalents directly to data obtained from cultured skin fibroblasts. Samples without paired germline tissue were analyzed using a combination of population allelic frequency (minor allele threshold of 0.01% or lower), VAF (with likely germline VAFs considered to be between 30 and 60% for genes on autosomal chromosomes and 80% or higher for genes on the X chromosome), and the frequency of the variant in question in tumor databases such as COSMIC. Any variant passing the above population filters, but which still occurred more than twice in COSMIC, was considered to not be of germline origin.

This filtering process produced a list of variants of likely somatic or definitive somatic origin which we reviewed manually for clinical and biological relevance. The determination of “likely somatic” or “somatic” origin adhered to criteria defined in the original RUNX1 database manuscript.^15^ “Clinically relevant” variants were known pathogenic germline variants in leukemia or variants that were present more than twice in COSMIC in hematopoietic and lymphoid samples (H&L samples). Novel driver variants were clinically relevant if they were present in a gene known to be recurrently mutated in COSMIC H&L samples, were a truncating variant (nonsense, frameshift indels, essential splice site variants), were in the same domain as known pathogenic variants (for example, the RUNT domain in RUNX1), or were a deletion in a gene where deletion is a known mechanism of disease. Missense variants were considered to be clinically relevant if they were damaging in at least 3 *in silico* algorithms and were highly conserved via GERP and Phylop scores. All somatic and likely somatic variants that did not meet criteria for clinical relevance were categorized as “possibly relevant” or “of unknown relevance”.

